# Appendage regeneration requires IMPDH2 and creates a sensitized environment for enzyme filament formation

**DOI:** 10.1101/2024.07.29.605679

**Authors:** Morgan E. McCartney, Gavin M. Wheeler, Audrey G. O’Neill, Jeet H. Patel, Zoey R. Litt, S. John Calise, Justin M. Kollman, Andrea E. Wills

## Abstract

Regeneration of lost tissue requires biosynthesis of metabolites needed for cell proliferation and growth. Among these are the critical purine nucleotides ATP and GTP. The abundance and balance of these purines is regulated by inosine monophosphate dehydrogenase 2 (IMPDH2), which catalyzes the committing step of GTP synthesis. IMPDH2 assembles into filaments that resist allosteric inhibition under conditions of high GTP demand. Here we asked whether IMPDH2 is required in the highly proliferative context of regeneration, and whether its assembly into filaments takes place in regenerating tissue. We find that inhibition of IMPDH2 leads to impaired tail regeneration and reduced cell proliferation in the tadpole *Xenopus tropicalis*. We find that both endogenous and fluorescent fusions of IMPDH2 robustly assemble into filaments throughout the tadpole tail, and that the regenerating tail creates a sensitized condition for filament formation. These findings clarify the role of purine biosynthesis in regeneration and reveal that IMPDH2 enzyme filament formation is a biologically relevant mechanism of regulation in vertebrate regeneration.

**Graphical Abstract:** 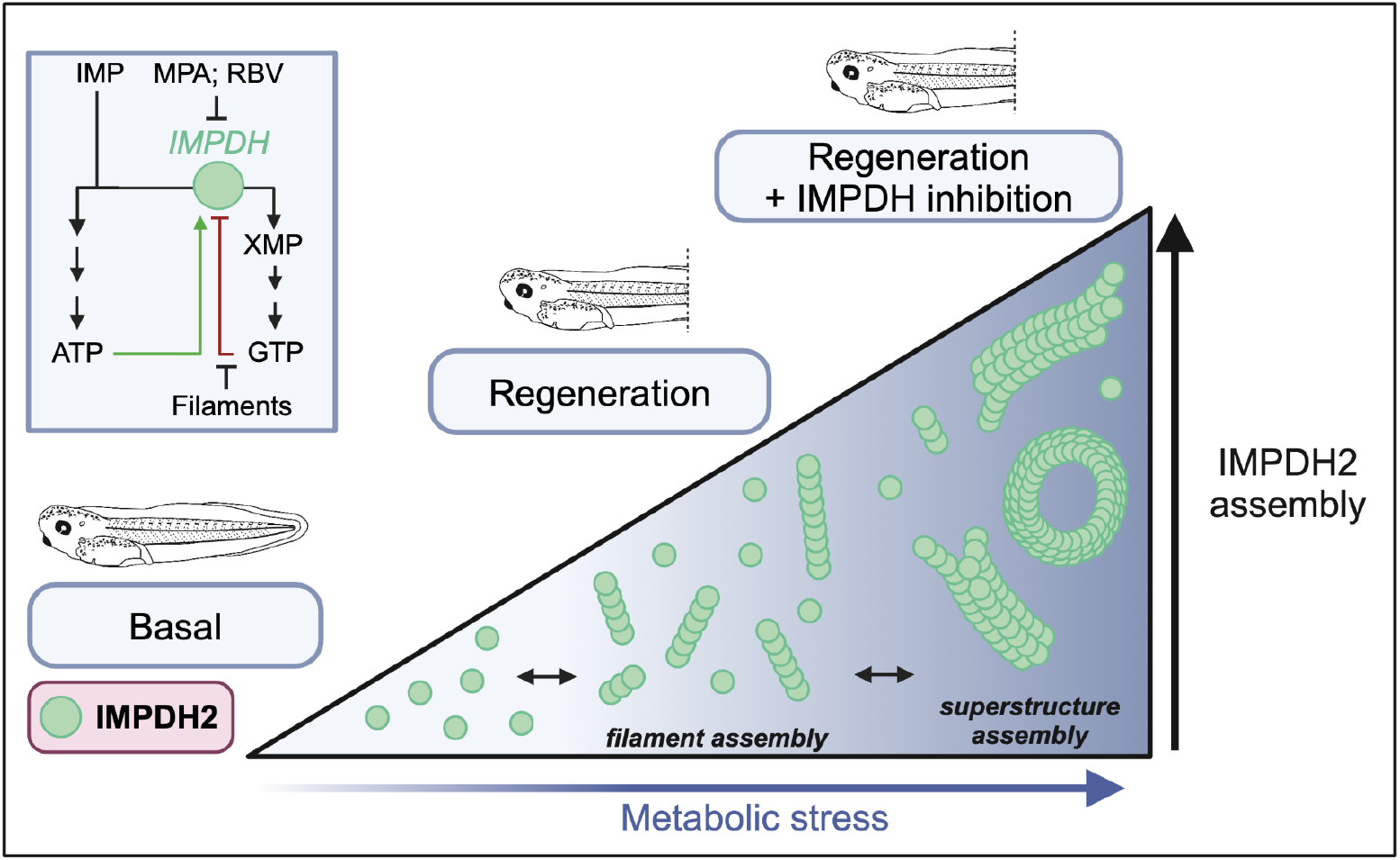

## Introduction

The event of a major injury creates a demand for biomolecules used in the synthesis of new cells to replace lost or damaged tissue. Our fundamental hypothesis is that after injury, this demand is met by reprogramming aspects of metabolism to funnel metabolites toward biosynthesis. Recently, we described the pentose phosphate pathway (PPP) as a principal fate for glucose in proliferating cells during regeneration.^1^ The PPP yields several useful intermediates for biosynthesis, especially purine nucleotides.^2^ During cell proliferation, nucleotides are in high demand for DNA and RNA synthesis, cytoskeletal remodeling, and energy. Following up on those observations, we turned our attention here to the regulation of nucleotide biosynthesis during tail regeneration. Downstream of the PPP, the enzyme inosine monophosphate dehydrogenase (IMPDH) sits at a major regulatory node in *de novo* nucleotide biosynthesis by catalyzing the committing step in the production of GTP rather than ATP^3^ (**Figure 1A**). Structurally, IMPDH constitutively forms tetramers that reversibly dimerize into either extended, active octamers in the presence of ATP or compressed, inactive octamers in the presence of GTP.^4^ Under conditions of purine stress, IMPDH octamers assemble into filaments that structurally resist GTP-induced octamer compression, sustaining enzyme activity in the presence of normally inhibitory levels of GTP.^4^ Filaments can further assemble laterally to create filament superstructures that can be several microns long.^5–9^ The role of IMPDH, and the physiological relevance of filament formation, has not been tested in a regenerative context.

**Figure 1.**
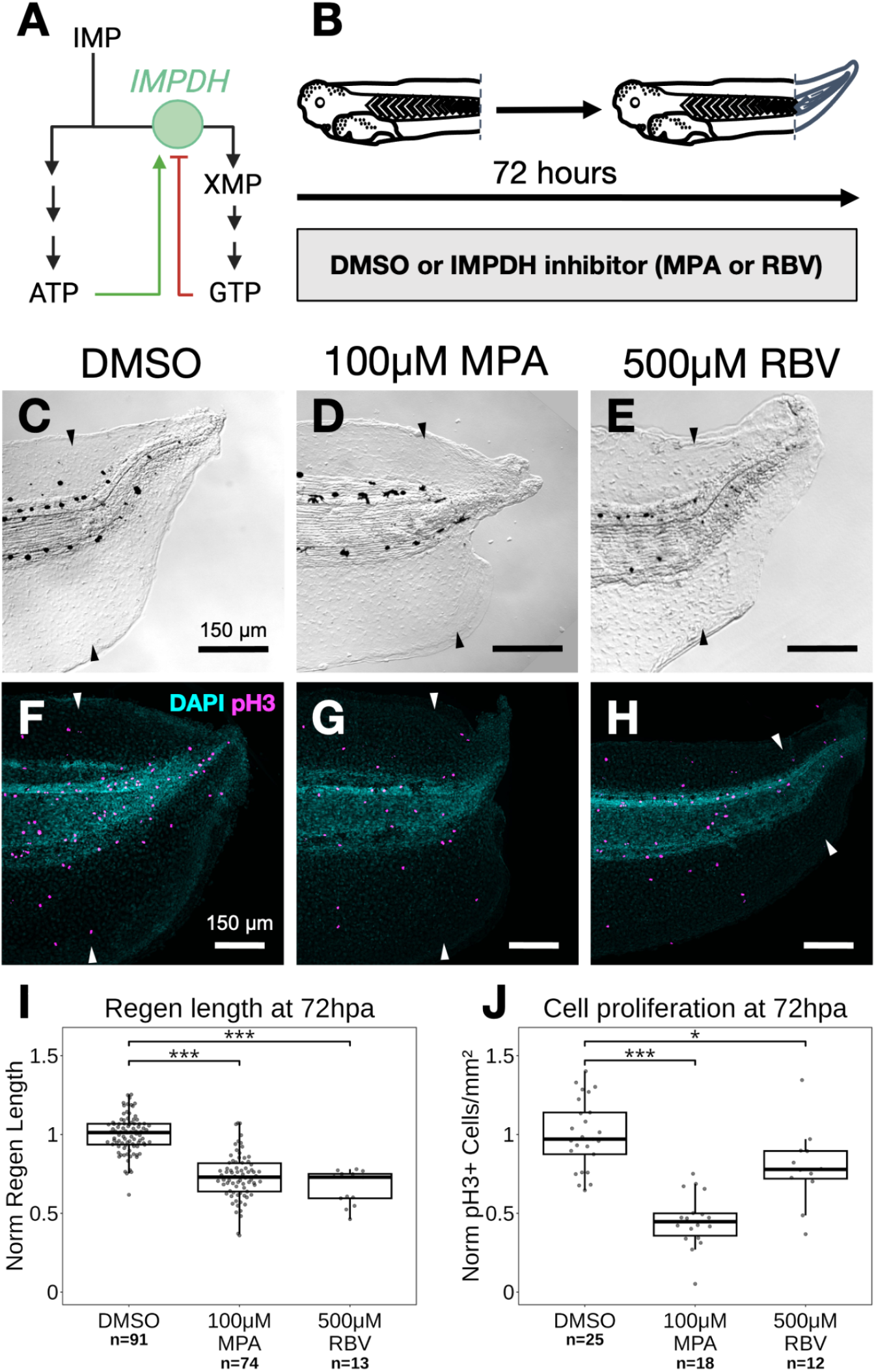
IMPDH inhibition impairs regeneration and cell proliferation. (A) Simplified pathway diagram of IMPDH2 highlighting regulation by purine nucleotides. (B) Schematic of experimental design. Tadpoles are amputated and then allowed to regenerate for 72 hours in media supplemented with either vehicle control (DMSO) or IMPDH inhibitor (MPA or RBV). (C-E) DIC images of tadpole tails 72 hours post amputation (hpa) treated with either 0.05% DMSO, 100 μM MPA, or 500 μM RBV. Arrows indicate the plane of amputation. (F-H) Immunohistochemistry (IHC) for proliferative marker pH3 at 72 hpa. Arrows indicate the plane of amputation. (I-J) Quantification of regeneration defects following IMPDH inhibition. Regenerate length (I) and pH3^+^ cell density in regenerating tissue (J) normalized to DMSO condition by experimental clutch at 72 hpa. Statistical significance between conditions was determined by ANOVA followed by Tukey’s HSD test. *p < 0.05; *** p <0.001.

## Results and Discussion

### IMPDH2 is required for cell proliferation and growth during tail regeneration

We first hypothesized that IMPDH2, the isoform of IMPDH commonly expressed in proliferating cells^10–13^, might play a specific role in regenerating tissues with high purine demand. To ask if IMPDH2 was required during regeneration, we amputated tadpole tails and treated with the noncompetitive IMPDH2 inhibitor mycophenolic acid (MPA)^14–17^ (**Figure 1B**). As an alternative inhibitor, we also used the competitive inhibitor Ribavirin (RBV)^18^. Treatment during regeneration with either 100 µM MPA or 500 µM RBV resulted in significantly shorter regenerated tails at 72 hours post amputation (hpa) (Figure **1C-E, I**). Consistent with previous reports that cell proliferation is sensitive to IMPDH2 loss^10–13^, we detected a significant reduction in mitotic nuclei in MPA or RBV treated regenerating tadpoles (**Figure 1F-H, J**). Thus, inhibition of IMPDH2 impairs normal tail regeneration and limits cell proliferation in *Xenopus* tadpoles. A remaining question is which of the many possible functions of a carefully regulated balance of ATP and GTP are most sensitive to IMPDH2 loss. Among the most expensive cell functions for GTP during proliferation are DNA synthesis, protein synthesis, and spindle assembly, as each tubulin monomer requires GTP hydrolysis for microtubule assembly.^19^ We conjecture that these functions would rely heavily on IMPDH to maintain sufficient levels of GTP relative to ATP.

### IMPDH2 expression during tail regeneration

Our next hypothesis was that IMPDH2 might be strongly upregulated during regeneration to meet the increasing purine demand of cell proliferation. We were surprised to find that in our previously published bulk RNA-Seq data^20^, most purine biosynthesis enzymes are significantly upregulated at 24 hpa, but IMPDH2 itself is not significantly changed (**Figure 2A, B**). To look more closely at IMPDH2 expression and localization at other regenerative timepoints, we used *in situ* hybridization. Prior to injury, IMPDH2 is expressed at low levels in the tail tip (**Figure 2C**). During regeneration, we saw generally unchanged expression except at 72 hpa, when expression was upregulated in the tail tip. 72 hpa coincides with the time at which cell proliferation increases during regeneration.^21,22^ However, protein levels of IMPDH2 are comparable in regenerating tails at 72hpa relative to uninjured stage matched tails, and actually decrease markedly in both uninjured and regenerating tadpoles during the timecourse of regeneration (**Figure 2D, Supplemental Figure 1**).

**Figure 2.**
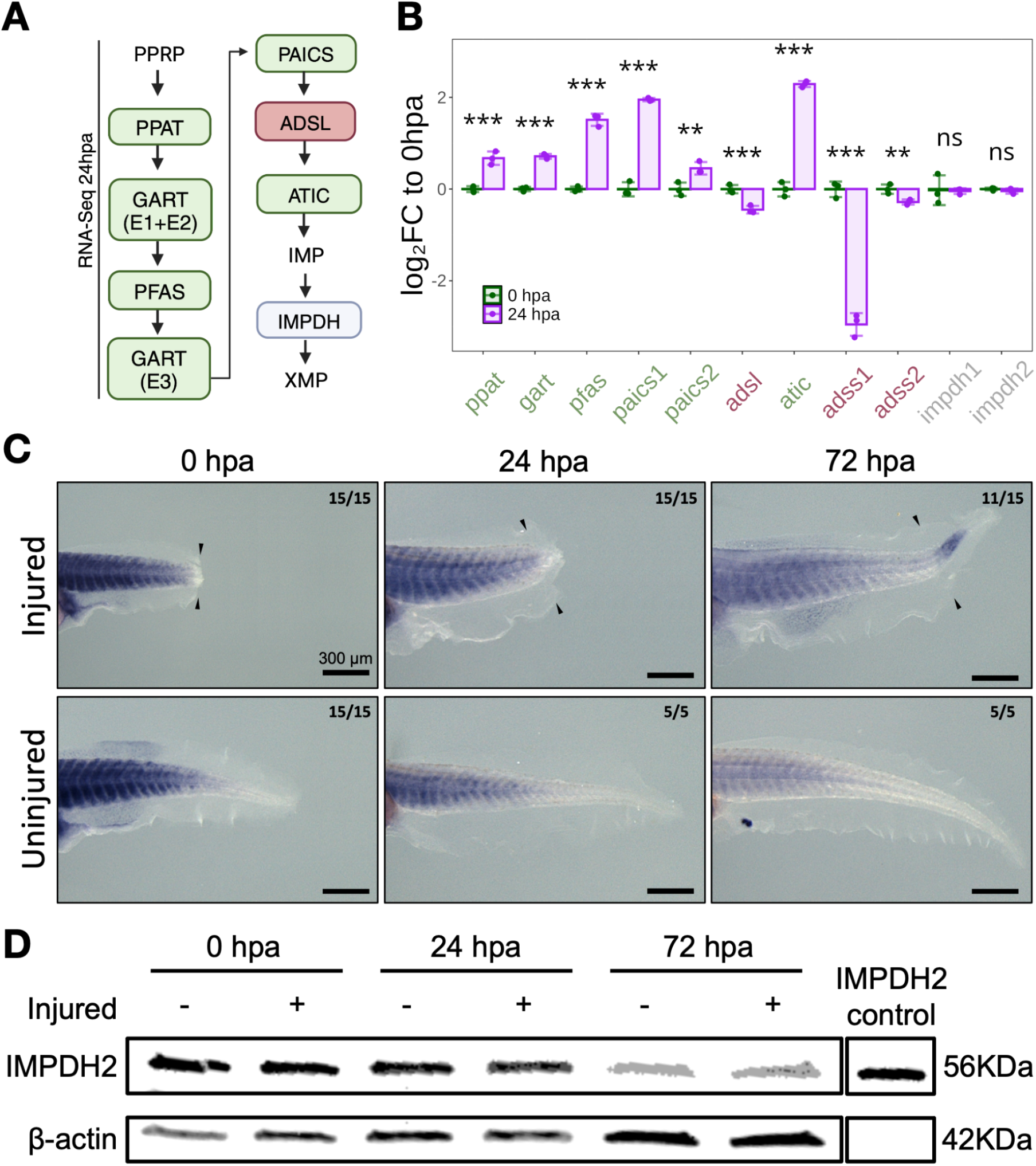
IMPDH2 expression during regeneration. (A) Schematic of the *de novo* purine synthesis pathway upstream of IMPDH. Phosphoribosyl pyrophosphate (PRPP) enters and is converted into inosine monophosphate (IMP) before commitment by IMPDH into xanthine monophosphate (XMP) and ultimately GTP. Enzymes are boxed with fill color representing an overall increase (green) or decrease (red) in transcript levels at 24 hpa. (B) Expression of *de novo* purine synthesis genes at 0 and 24 hpa as log_2_FC to 0 hpa mean. Statistical significance was determined using EdgeR. **p < 0.01; ***p < 0.001; ns, not significant. (C) *In situ* hybridization of *impdh2* at 0, 24, and 72 hpa in injured and uninjured tadpoles. Expression appears concentrated in muscle tissue, with somitic boundaries clearly visible. Overall expression declines during development, but increases specifically in the regenerating tissue at 72 hpa. Numbers in the top right of each panel indicate the number of samples that resemble the representative image over the total number assayed. Arrows indicate the plane of amputation. (D) Western blot for IMPDH2 and β-actin in injured (+) or uninjured (-) tails at 0, 24, and 72 hpa.

### IMPDH2 forms filaments in the tail following inhibition

The modest changes in IMPDH2 expression led us to wonder if other mechanisms of IMPDH2 regulation might contribute to its function during regeneration. Specifically, allosteric regulation by filament formation might be a different mode by which IMPDH2 activity is modulated in this context. Under conditions of purine stress (such as IMPDH2 inhibition), IMPDH2 forms filaments and laterally-bundled filament superstructures (sometimes called “rods and rings” or cytoophidia) that allow the enzyme to maintain activity by preventing feedback inhibition.^4^ We therefore queried whether IMPDH2 forms filaments in uninjured or regenerating tadpoles.
IMPDH2 was diffuse throughout the uninjured tadpole tail (**Figure 3A, B, E**). However, after treatment with MPA or RBV, filamentous superstructures formed throughout the tail (**MPA: Figure 3C, F; RBV: Figure 3D, G**). These superstructures were most abundant in axial tissues, where they were distributed in a segmental pattern that matched the periodicity of somites (**Figure 3C, D**). To observe superstructures in live tissue, we injected mRNA encoding an IMPDH2-RFP fusion protein at the 2 cell stage. These embryos were reared to stage 41, at which point they were treated with MPA and live imaged (**Figure 3H**), which also induced filament superstructures (**Figure 3I, J**).

**Figure 3.**
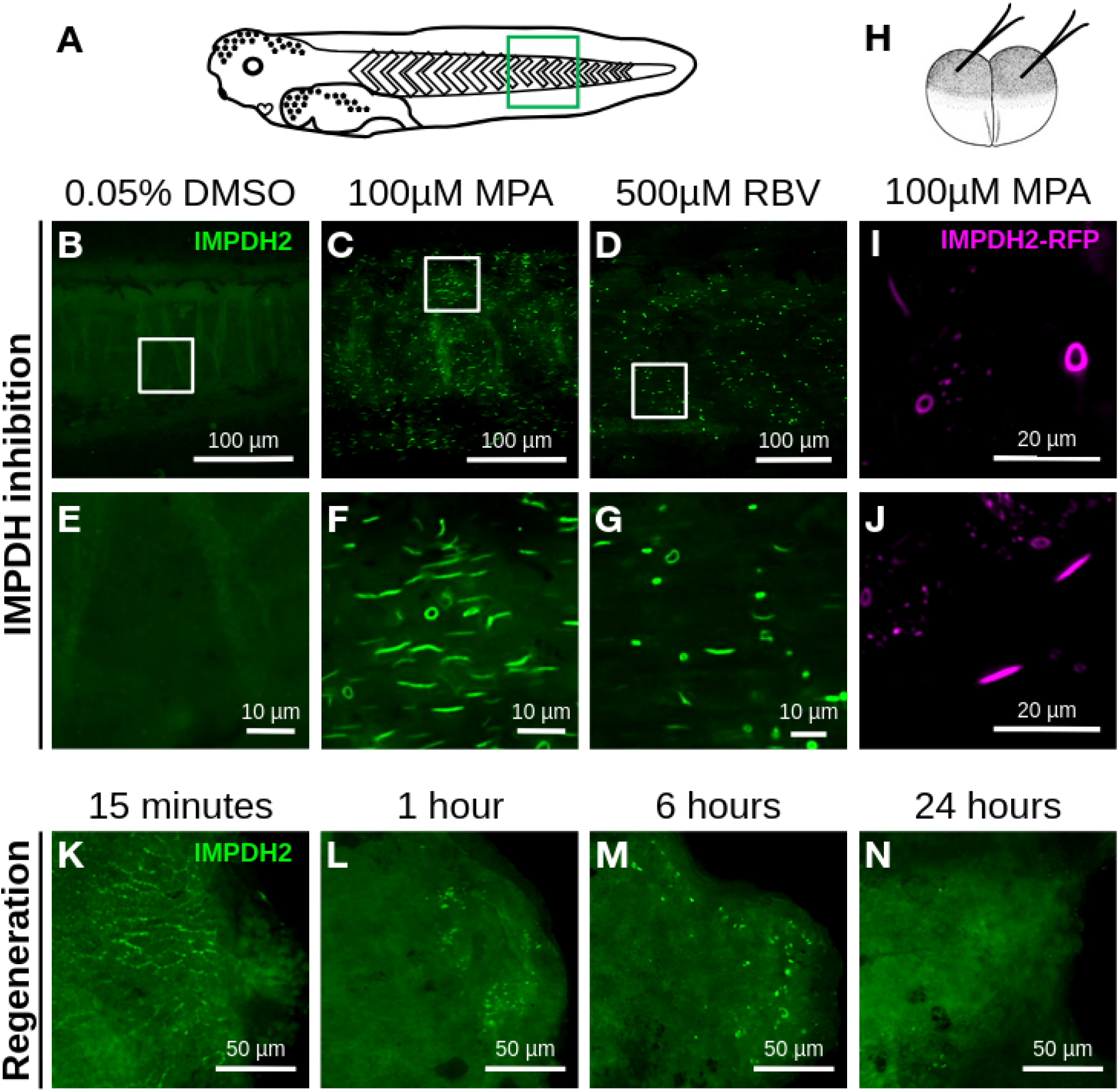
IMPDH2 forms filaments in the tail following inhibition. (A) Tadpole diagram. Green box denotes approximate imaging location. (B-D) Immunohistochemistry (IHC) for IMPDH2 in St. 47 tadpole tails after 72 hour treatment with 0.05% DMSO (B), 100 µM MPA (C), or 500 µM RBV (D). White boxes indicate location of close-up images shown in (E)-(G). (E-G) Close-up images of panels (B), (C), and (D) respectively. (H) mRNA microinjection diagram at the 2-cell stage. Embryos were injected with 1000 pg of mRNA encoding IMPDH2-RFP fusion protein. (I-J) Images of live St. 45 tadpoles expressing IMPDH2-RFP fusion protein after 24 hour treatment with 100 µM MPA. IMPDH2-RFP forms both Ring Superstructures (I) and Rod Superstructures (J) in the tadpole tail under IMPDH2 inhibition. (K-N) IHC for IMPDH2 during wound closure and early regeneration. Images were taken of the axial tissue at the amputation site. (K) At 15 minutes post amputation, IMPDH2 appears localized to the membrane in epidermal cells proximal to the amputation site in 7/8 samples. (L) By 1 hour small punctae begin to form near the injury in 6/7 samples. (M) By 6 hours the small punctae appear to have assembled into slightly larger structures reminiscent of “rod and ring” superstructures in 9/10 samples. (N) By 24 hours the signal around the wound site is once again diffuse and mostly free of any structures in 8/10 samples.

It is clear that *Xenopus* are able to assemble both endogenous and overexpressed IMPDH2 into filament superstructures when IMPDH2 is inhibited. It is intriguing that these were especially abundant in the somites. This coincides with the stronger somitic expression of *impdh2*, which is also seen in zebrafish^23^ (**Figure 2C**). This raises questions about potential cell type-specific functions for IMPDH2 and filament assembly. Notably, other enzymes in the purine biosynthesis pathway, specifically adenylosuccinate lyase (ADSL), are required for muscle development in *Xenopus*.^24^ Patient alleles of IMPDH2 that affect filament formation are associated with motor deficits, which may be attributable to changes in purinogenesis in either muscle cells, neurons, or both.^3^ As both muscle and neurons are cell types with high dynamic demand for ATP and GTP in cytoskeletal roles, these observations raise the possibility that IMPDH2 may play specific roles in these cell types during development.

### The regenerating tail creates a sensitized environment for IMPDH2 filament formation

We next hypothesized that regenerating tissues would have a greater purine demand than uninjured tissue, and thus be more likely to form IMPDH2 filaments. We therefore examined IMPDH2 localization during a regenerative timecourse, in the absence of inhibitor. Shortly after amputation, IMPDH2 transiently localized to cell membranes and punctae near the amputation plane (**Figure 3K-N**). Later in regeneration, however, filamentous structures were rarely observed (**Figure 4A-D**). The rapid early dynamics of IMPDH2 punctae after injury suggest to us that there may be rapid shifts in the localization of IMPDH2 to meet purine needs. A number of tissues show context-specific or dynamic IMPDH2 localization during their development, disease, or aging, including the retina and the developing mouse embryo.^25–28^ We also note that the filament superstructures that we observe following MPA treatment are large, micron-scale bundles of laterally associated filaments. The smaller punctae we observe immediately after injury may represent smaller bundles, but small individual filaments or multi-enzyme assemblies like purinosomes^29^ may also form during regeneration.

**Figure 4.**
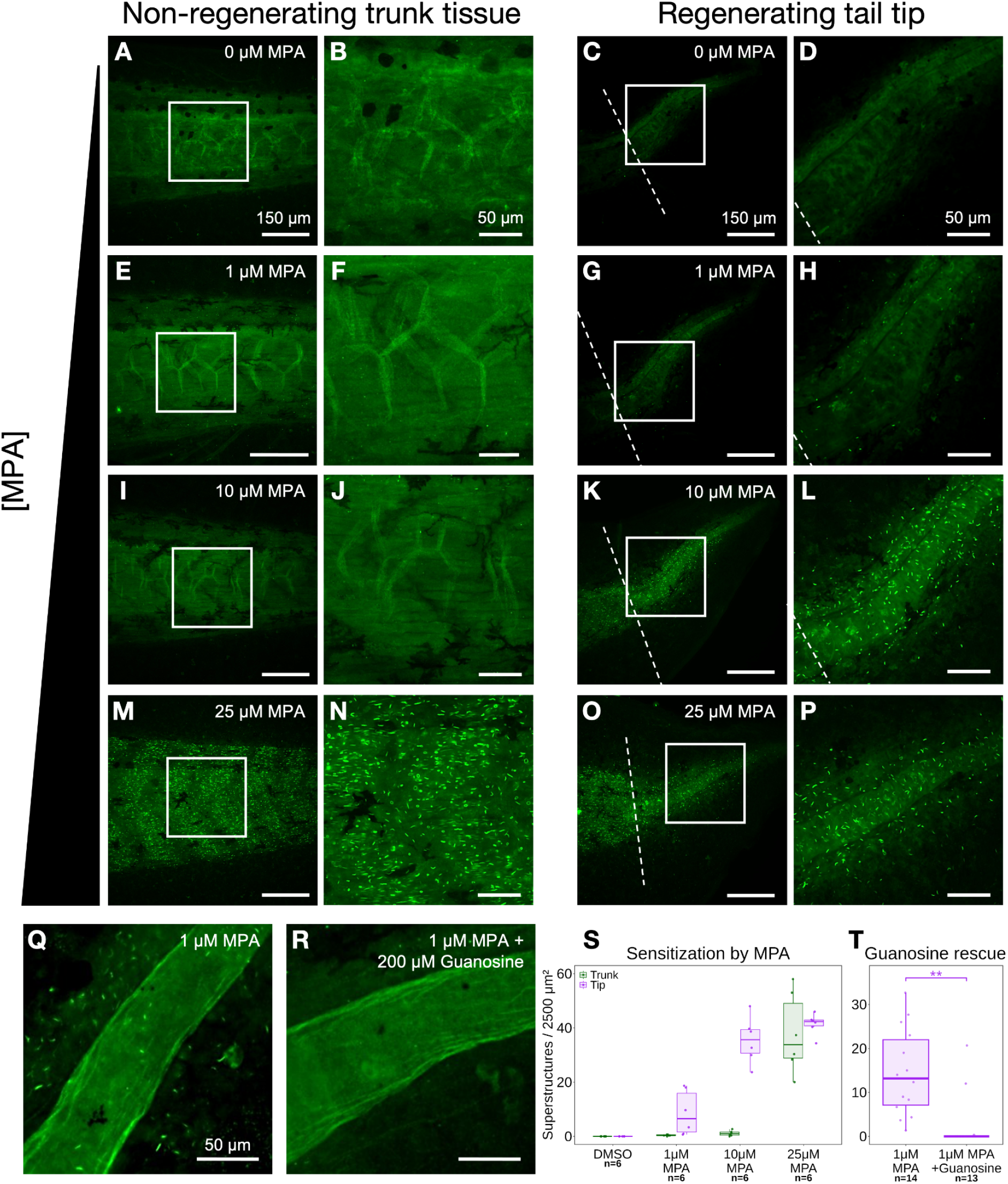
The regenerating tail creates a sensitized environment for IMPDH2 filament formation. (A-D) IHC for IMPDH2 in 72 hpa tadpoles treated with 0.05% DMSO. IMPDH2 filaments rarely assemble under control conditions in the non-regenerating trunk tissue (A and B) or the regenerating tail tip (C and D). (E-H) IHC for IMPDH2 in 72 hpa tadpoles treated with 1 µM MPA. IMPDH2 filaments do not assemble in the non-regenerating trunk tissue (E and F). Filaments do assemble in the regenerating tail tissue (G and H). (I-L) IHC for IMPDH2 in 72 hpa tadpoles treated with 10 µM MPA. IMPDH2 forms few filaments in the non-regenerating trunk tissue (I and J). IMPDH2 filaments assemble robustly in the regenerating tail tip (K and L). (M-P) IHC for IMPDH2 in 72 hpa tadpoles treated with 25 µM MPA. IMPDH2 filaments assemble robustly in the non-regenerating trunk tissue (M-N) and the regenerating tail tip (O-P). (Q-R) IHC for IMPDH2 in 72 hpa tadpoles. IMPDH2 filaments form in the regenerating tail tip after treatment with 1 µM MPA (Q). Filament formation is reduced in the regenerating tail tip after treatment with 1 µM MPA and 200 µM guanosine (R). (S) Quantification of IMPDH2 superstructures per 2500 µm^2^ in either the non-regenerating trunk tissue or the regenerating tail tip treated with either 0.05% DMSO, 1 µM MPA, 10 µM MPA, or 25 µM MPA. (T) Quantification of IMPDH2 superstructures per 2500 µm^2^ in tadpole tail tips treated with 1uM MPA vs 1 µM MPA + 200 µM guanosine. Statistical significance was determined by Welch’s t-test. **p<0.01.

Although we did not observe filaments in later stages of regeneration in the absence of inhibitor, we considered that regenerating tissue might be close to the threshold of filament formation. It would therefore be sensitized to form superstructures by doses of inhibitor that are too low to induce superstructures in non-regenerating tissue. Consistent with this hypothesis, we found that treatment with low doses of MPA (1-10 µM) induced abundant superstructures in regenerating tail tips, but not in uninjured anterior trunk tissue (**Figure 4E, F, I, J** vs **Figure 4G, H, K, L**), a significant difference (**Figure 4S**). Higher doses of MPA (25 µM) induced superstructures in all tissues (**Figure 4M-P**). A similar sensitization was seen for RBV (**Supplementary Figure 2**). To confirm that superstructures are formed in the sensitized condition due to purine depletion, we rescued filament formation by supplying guanosine as an alternative source of GTP. Guanosine is converted into guanine and subsequently GMP by the purine salvage pathway, bypassing IMPDH2 in the *de novo* synthesis pathway. We found that supplementation with 200 µM guanosine was sufficient to rescue the assembly of superstructures in regenerating tadpoles treated with 1 µM MPA (**Figure 4Q vs R, T**).

The sensitization of regenerating tissue to filament formation suggests that the demand for purines, or GTP specifically, is higher in regenerating tissue than in similar non-regenerating tissue. This is consistent with purine biosynthesis being one potential target of the observed upregulation of the PPP in regeneration^1^, although others may well contribute, such as NADPH production for redox homeostasis or lipid biosynthesis, as has also been suggested.^30,31^ Overall, the propensity for IMPDH2 filament assembly represents a useful insight toward understanding how injured tissue mobilizes its metabolic resources toward wound healing and cell proliferation: if ATP and GTP are sunk into DNA or RNA biosynthesis, both of these purines would need to be replenished. Derepression of IMPDH2 by filament formation would also allow GTP to be rapidly produced under cell conditions where it is specifically needed in abundance.

## Methods

### Xenopus tropicalis husbandry and use

Use of *Xenopus tropicalis* was carried out under the approval and oversight of the IACUC committee at UW, an AALAC-accredited institution, under animal protocol 4374-01. Ovulation of adult *Xenopus tropicalis* and generation of embryos by natural matings were performed according to published methods.^32,33^ Embryos were reared as described in Khokha et al.^32^ Staging was assessed by the Nieuwkoop and Faber (NF) staging series.^34^ Tadpoles were reared at 22 °C to stage 41 (3 dpf) and experiments were performed as described below. Animals were reared in petri dishes at a density of no more than 2 tadpoles/mL of rearing media, 1/9th Modified Ringer’s solution (1/9x MR), and clutchmates were randomly assigned to treatment groups. Sex is not able to be determined by this stage and so is not a relevant biological variable. Tadpoles do not begin to feed independently until stage 45/46 and were not fed during the course of these experiments.

### Xenopus tropicalis amputation assay

NF stage 41 tadpoles were anesthetized with 0.016% MS-222 in 1/9x MR and tested for response to touch prior to amputation surgery. Once fully anesthetized, a sterilized scalpel was used to amputate the posterior third of the tail. Amputated tadpoles were removed from anesthetic media within 10 minutes of amputation into 1/9x MR with or without added pharmacological inhibitors. Tadpoles were kept at a density of no more than 2.5 tadpoles per mL. For measurements of IMPDH2 inhibition and sensitized superstructure assembly, tadpoles were fixed at 72 hpa. For observation of early regeneration, tadpoles were fixed at 15 minutes, 1 hour, 6 hours, and 24 hours post amputation. Tadpoles were fixed in 1x MEM with 3.7% formaldehyde for 50 minutes at room temperature.

### Pharmacological inhibition

Mycophenolic Acid (MPA) (MilliporeSigma M5255) was resuspended to a 200 mM stock in DMSO and Ribavirin (RBV) (Cayman Chemical 16757) to a 500 mM stock in DMSO. Guanosine (MilliporeSigma G6264) was resuspended directly in 1/9x MR to a concentration of 200 μM immediately prior to amputation. To verify IMPDH2 superstructure formation in tadpoles, uninjured and injured tadpoles were reared with 0.05% DMSO, 100 µM MPA, or 500 µM RBV diluted in 1/9x MR until collection at 72 hours following treatment. To induce a sensitized-regenerative condition, injured tadpoles were reared with 0.05% DMSO, 1 µM MPA, 10 µM MPA, 25 µM MPA, 200 µM Guanosine, or 1 µM MPA + 200 µM Guanosine diluted in 1/9x MR until collection at 72 hours following treatment. RBV was also used to characterize the sensitized-regenerative condition and is reported in **Supplementary Figure 2**.

### Tail regeneration length analysis

Stereoscope imaging for regeneration length quantification was performed on a Leica M205 FA with a color camera. Fixed tadpoles were imaged in 1x PBS on 1% agarose pads and measurements were recorded using LAS X software. Representative images were acquired using a Leica DM 5500 B microscope using a 10x objective and processed using FIJI image analysis software.^35^

### Immunohistochemistry

Fixed tadpoles were permeabilized by washing 3 × 20 minutes in 1x PBS + 0.01% Triton X-100 (PBS-Triton). Tadpoles were blocked for 1 hour at room temperature in 10% CAS-block (Invitrogen #00-8120) in PBS-Triton. Tadpoles were then incubated in primary antibody [1:100 rabbit anti-IMPDH2, proteintech 12948-1-AP; 1:1000 mouse anti-Histone H3 (phospho S10), Abcam ab14955] diluted in 100% CAS-block overnight at 4°C. Tadpoles were washed 3 × 10 minutes at room temperature in PBS-Triton then blocked for 30 minutes in 10% CAS-block in PBS-Triton. Secondary antibody (goat anti-rabbit 488, Invitrogen A11008; goat-anti mouse 594, Abcam ab150116) was diluted 1:500 in 100% CAS-block and incubated for 2 hours at room temperature. Tadpoles were then washed 3 × 10 minutes in PBS-Triton followed by a 10 minute incubation in 1:2000 DAPI (Sigma D9542) in PBS-Triton before 3 × 20 minute washes in PBS-Triton. Isolated tails were mounted on slides in ProLong Diamond (ThermoFisher P36970). Images were acquired using a Leica DM 5500 B microscope with 10X, 20X and 40X objectives and processed using Fiji image analysis software.

### Cell proliferation assay

Images of pH3 stains were imported into Fiji. The amputation plane was identified, and the regenerating tissue area was measured. Then, the number of pH3 positive cells within the regenerating tissue was counted. Density of pH3 cells was determined by dividing the number of pH3 positive cells by the total area of regenerating tissue. Density was normalized to DMSO controls so that data from multiple clutches could be analyzed in tandem.

### Whole-mount *in situ* hybridization

Tadpoles were fixed in 1x MEM with 3.7% formaldehyde for 45 minutes at room temperature. *Xenopus tropicalis* multibasket *in situ* hybridization protocols were followed as described by Khokha et al.^32^, with pre-hybridization performed overnight. Whole tadpoles were imaged in 1x PBS + 0.1% Tween-20 on a bed of 1% agarose using a Leica M205 FA stereomicroscope with a color camera. Probes were synthesized using the following primer pair designed against a single exon of the *impdh2* transcript: forward - gcaagttgcccattgttaatggc, reverse - TAATACGACTCACTATAGGGaaggaccacggcatccac.

### Western blotting

Tadpoles were anesthetized with 0.016% MS-222 in 1/9x MR and tested for response to touch prior to amputation surgery. For each experimental group, 10 tadpole tails were amputated just posterior to the vent and homogenized on ice in 100 μL of lysis buffer (50 mM Tris pH 7.6, 150 mM NaCl, 10 mM EDTA, 0.1% Triton X-100, Roche cOmplete^TM^ Protease Inhibitor Cocktail). Homogenized samples were centrifuged at 18,400xg for 20 minutes at 4°C. The soluble fraction was collected, and total protein concentration was quantified with a Pierce BCA assay (Thermo Scientific 23227). Samples were denatured by adding 1/4 volume of 4X Protein Sample Loading Buffer (Licor 928-40004). The total protein concentration of each sample was normalized according to the BCA assay by adding additional 1X Protein Sample Loading Buffer (4X Protein Sample Loading Buffer diluted to 1X with lysis buffer). Samples were heated at 100 °C for 5 minutes. Equal inputs of the samples were run on a 4-20% Mini-PROTEAN TGX precast gel (BIO-RAD 4561094) at 180 V for 40 minutes in manufacturer recommended running buffer (25 mM Tris, 192 mM glycine, 0.1% SDS, pH 8.3). Transfer to a nitrocellulose membrane was done using an Invitrogen Power Blotter Select Transfer Stack (Thermo Fisher PB3310) on an Invitrogen Power Blotter System (Thermo Fisher PB0012) using the Mixed Range MW Pre-Programmed method (constant 2.5 A with 25 V limit for 7 minutes). Membrane was incubated in Intercept PBS Blocking Buffer (Licor 927-70001) for 1 hour at 25°C with rocking. Membrane was added into 5 mL of 1X PBS-Tween (137 mM NaCl, 2.7 mM KCl, 10 mM Na2HPO4, 1.8 mM KH2PO4, 0.1% w/v Tween-20 detergent) and 5 mL of Intercept PBS Blocking Buffer. Membrane was incubated in primary antibodies: 1:1000 rabbit anti-IMPDH2 (Proteintech 12948-1-AP) and 1:3000 mouse anti-β-actin (Santa Cruz Biotech sc-47778) overnight, shaking at 4 °C. Membrane was washed with 1X PBS-Tween for 4x5min, then added into 5mL of 1X PBS-Tween mixed with 5mL of Intercept PBS Blocking Buffer. Membrane was incubated in the dark for 1 hour, shaking at 25 °C with secondary antibodies: 1:10000 goat anti-rabbit IgG (H+L) (DyLight 800 Fisher PISA510036) and 1:10000 goat anti-mouse IgG (H+L) (DyLight 680 Fisher PI35518). After 4 x 5 min washes in 1X PBS-Tween, the membrane was imaged on a LI-COR Imaging System (LI-COR Biosciences). Quantification of western blots was done in Fiji.^35^

### mRNA synthesis and injections

Plasmid pCS107_IMPDH2-RFP was synthesized using Gibson assembly following PCR amplification with the following primers: hIMPDH2 template (forward - GCTCGCCACCatggccgactacctgattagtg, reverse - GTACCGTCGAgaaaagccgcttctcatacgaa) and pCS107_Cterm-RFP (forward - gcggcttttcTCGACGGTACCGCGGGCCCG, reverse - tagtcggccatGGTGGCGAGCTCGAGATCCTG). Final construct was sequence verified before *in vitro* transcription. Plasmid DNA was linearized using *AscI* enzyme and mRNA was transcribed with SP6 mMessage mMachine kit (Invitrogen). mRNA was injected into embryos at the 2-cell stage at a dose of 1000 pg/embryo.

### Live imaging

Tadpoles were anesthetized with 0.016% MS-222 in 1/9x MR and then mounted within 1% LMP agarose on coverslip-bottom dishes as described by Kieserman et al.^36^ Immobilized tadpoles were imaged using a Leica SP8 confocal microscope with a 20x objective.

### Analysis of previously published datasets

Published RNA-sequencing data (CPM and statistical analysis) was obtained from the NCBI Gene expression Omnibus (GEO) under accession number GSE174798.^20^

### Quantification and statistical analysis

Boxplots were generated using the R package ggplot2.^37^ Length and cell proliferation measurements were compared using ANOVA and post hoc Tukey’s HSD test to identify differences between groups. For bulk RNA-seq analysis, statistically significant changes in gene expression were determined using edgeR.^38^ Superstructure density in the regenerating tails of tadpoles treated with MPA with or without guanosine was compared using Welch’s t-test. All statistical analyses were performed with R statistical software.^39^ Information regarding statistics can be found in the figure legends.

## Acknowledgements

This work was supported by a University of Washington Levinson fellowship to MEM, by a Helen Hay Whitney Foundation fellowship to SJC, and by grants R35GM149542 to JMK, R01GM148490 to AEW, and R01NS099124 to AEW. The authors gratefully acknowledge members of the Wills and Kollman labs for helpful discussion throughout the execution of this work. The graphical abstract and Figure 2A were generated using Biorender.

## Author contributions

Study conception and experimental design: MEM, GMW, AGO, JHP, JMK, AEW. Performing experiments: MEM, GMW, AGO, JHP, ZRL. Analyzing experiments: MEM, GMW, AGO, JHP, ZRL, AEW. Manuscript draft: MEM, GMW, AGO, AEW. Manuscript revision: MEM, GMW, AGO, JHP, ZRL, SJC, JMK, AEW. Funding: JMK, AEW.

## References

1. Patel, J.H., Ong, D.J., Williams, C.R., Callies, L.K., and Wills, A.E. (2022). Elevated pentose phosphate pathway flux supports appendage regeneration. Cell Rep. 41, 111552. 10.1016/j.celrep.2022.111552.

2. TeSlaa, T., Ralser, M., Fan, J., and Rabinowitz, J.D. (2023). The pentose phosphate pathway in health and disease. Nat. Metab. 5, 1275–1289. 10.1038/s42255-023-00863-2.

3. Burrell, A.L., and Kollman, J.M. (2022). IMPDH dysregulation in disease: a mini review. Biochem. Soc. Trans. 50, 71–82. 10.1042/BST20210446.

4. Johnson, M.C., and Kollman, J.M. (2020). Cryo-EM structures demonstrate human IMPDH2 filament assembly tunes allosteric regulation. eLife 9, e53243. 10.7554/eLife.53243.

5. Thomas, E.C., Gunter, J.H., Webster, J.A., Schieber, N.L., Oorschot, V., Parton, R.G., and Whitehead, J.P. (2012). Different Characteristics and Nucleotide Binding Properties of Inosine Monophosphate Dehydrogenase (IMPDH) Isoforms. PLoS ONE 7, e51096. 10.1371/journal.pone.0051096.

6. Juda, P., Šmigová, J., Kováčik, L., Bártová, E., and Raška, I. (2014). Ultrastructure of Cytoplasmic and Nuclear Inosine-5’-Monophosphate Dehydrogenase 2 “Rods and Rings” Inclusions. J. Histochem. Cytochem. 62, 739–750. 10.1369/0022155414543853.

7. Ji, Y., Gu, J., Makhov, A.M., Griffith, J.D., and Mitchell, B.S. (2006). Regulation of the interaction of inosine monophosphate dehydrogenase with mycophenolic Acid by GTP. J. Biol. Chem. 281, 206–212. 10.1074/jbc.M507056200.

8. Carcamo, W.C., Satoh, M., Kasahara, H., Terada, N., Hamazaki, T., Chan, J.Y.F., Yao, B., Tamayo, S., Covini, G., von Mühlen, C.A., et al. (2011). Induction of cytoplasmic rods and rings structures by inhibition of the CTP and GTP synthetic pathway in mammalian cells. PloS One 6, e29690. 10.1371/journal.pone.0029690.

9. Calise, S.J., Carcamo, W.C., Krueger, C., Yin, J.D., Purich, D.L., and Chan, E.K.L. (2014). Glutamine deprivation initiates reversible assembly of mammalian rods and rings. Cell. Mol. Life Sci. CMLS 71, 2963–2973. 10.1007/s00018-014-1567-6.

10. Jackson, R.C., Weber, G., and Morris, H.P. (1975). IMP dehydrogenase, an enzyme linked with proliferation and malignancy. Nature 256, 331–333. 10.1038/256331a0.

11. Senda, M., and Natsumeda, Y. (1994). Tissue-differential expression of two distinct genes for human IMP dehydrogenase (E.C.1.1.1.205). Life Sci. 54, 1917–1926. 10.1016/0024-3205(94)90150-3.

12. Collart, F.R., Chubb, C.B., Mirkin, B.L., and Huberman, E. (1992). Increased inosine-5’-phosphate dehydrogenase gene expression in solid tumor tissues and tumor cell lines. Cancer Res. 52, 5826–5828. 10.2172/10148922.

13. Nagai, M., Natsumeda, Y., and Weber, G. (1992). Proliferation-linked regulation of type II IMP dehydrogenase gene in human normal lymphocytes and HL-60 leukemic cells. Cancer Res. 52, 258–261.

14. Hedstrom, L. (1999). IMP dehydrogenase: mechanism of action and inhibition. Curr. Med. Chem. 6, 545–560.

15. Sintchak, M.D., Fleming, M.A., Futer, O., Raybuck, S.A., Chambers, S.P., Caron, P.R., Murcko, M.A., and Wilson, K.P. (1996). Structure and mechanism of inosine monophosphate dehydrogenase in complex with the immunosuppressant mycophenolic acid. Cell 85, 921–930. 10.1016/s0092-8674(00)81275-1.

16. Franklin, T.J., and Cook, J.M. (1969). The inhibition of nucleic acid synthesis by mycophenolic acid. Biochem. J. 113, 515–524. 10.1042/bj1130515.

17. Hager, P.W., Collart, F.R., Huberman, E., and Mitchell, B.S. (1995). Recombinant human inosine monophosphate dehydrogenase type I and type II proteins: Purification and characterization of inhibitor binding. Biochem. Pharmacol. 49, 1323–1329. 10.1016/0006-2952(95)00026-V.

18. Streeter, D.G., Witkowski, J.T., Khare, G.P., Sidwell, R.W., Bauer, R.J., Robins, R.K., and Simon, L.N. (1973). Mechanism of action of 1--D-ribofuranosyl-1,2,4-triazole-3-carboxamide (Virazole), a new broad-spectrum antiviral agent. Proc. Natl. Acad. Sci. U. S. A. 70, 1174–1178. 10.1073/pnas.70.4.1174.

19. Barisic, M., Rajendraprasad, G., and Steblyanko, Y. (2021). The metaphase spindle at steady state - Mechanism and functions of microtubule poleward flux. Semin. Cell Dev. Biol. 117, 99–117. 10.1016/j.semcdb.2021.05.016.

20. Patel, J.H., Schattinger, P.A., Takayoshi, E.E., and Wills, A.E. (2022). Hif1α and Wnt are required for posterior gene expression during Xenopus tropicalis tail regeneration. Dev. Biol. 483, 157–168. 10.1016/j.ydbio.2022.01.007.

21. Love, N.R., Chen, Y., Bonev, B., Gilchrist, M.J., Fairclough, L., Lea, R., Mohun, T.J., Paredes, R., Zeef, L.A.H., and Amaya, E. (2011). Genome-wide analysis of gene expression during Xenopus tropicalis tadpole tail regeneration. BMC Dev. Biol. 11, 70. 10.1186/1471-213X-11-70.

22. Kakebeen, A.D., Chitsazan, A.D., Williams, M.C., Saunders, L.M., and Wills, A.E. (2020). Chromatin accessibility dynamics and single cell RNA-Seq reveal new regulators of regeneration in neural progenitors. eLife 9, e52648. 10.7554/eLife.52648.

23. Wu, X., Zhong, H., Song, J., Damoiseaux, R., Yang, Z., and Lin, S. (2006). Mycophenolic acid is a potent inhibitor of angiogenesis. Arterioscler. Thromb. Vasc. Biol. 26, 2414–2416. 10.1161/01.ATV.0000238361.07225.fc.

24. Duperray, M., Hardet, F., Henriet, E., Saint-Marc, C., Boué-Grabot, E., Daignan-Fornier, B., Massé, K., and Pinson, B. (2023). Purine Biosynthesis Pathways Are Required for Myogenesis in Xenopus laevis. Cells 12, 2379. 10.3390/cells12192379.

25. Lake, J.I., Avetisyan, M., Zimmermann, A.G., and Heuckeroth, R.O. (2016). Neural crest requires Impdh2 for development of the enteric nervous system, great vessels, and craniofacial skeleton. Dev. Biol. 409, 152–165. 10.1016/j.ydbio.2015.11.004.

26. Cleghorn, W.M., Burrell, A.L., Giarmarco, M.M., Brock, D.C., Wang, Y., Chambers, Z.S., Du, J., Kollman, J.M., and Brockerhoff, S.E. (2022). A highly conserved zebrafish IMPDH retinal isoform produces the majority of guanine and forms dynamic protein filaments in photoreceptor cells. J. Biol. Chem. 298, 101441. 10.1016/j.jbc.2021.101441.

27. Plana-Bonamaisó, A., López-Begines, S., Fernández-Justel, D., Junza, A., Soler-Tapia, A., Andilla, J., Loza-Alvarez, P., Rosa, J.L., Miralles, E., Casals, I., et al. (2020). Post-translational regulation of retinal IMPDH1 in vivo to adjust GTP synthesis to illumination conditions. eLife 9, e56418. 10.7554/eLife.56418.

28. Calise, S.J., O’Neill, A.G., Burrell, A.L., Dickinson, M.S., Molffno, J., Clarke, C., Quispe, J., Sokolov, D., Buey, R.M., and Kollman, J.M. (2024). Light-sensitive phosphorylation regulates retinal IMPDH1 activity and filament assembly. J. Cell Biol. 223, e202310139. 10.1083/jcb.202310139.

29. Zhao, H., Chiaro, C.R., Zhang, L., Smith, P.B., Chan, C.Y., Pedley, A.M., Pugh, R.J., French, J.B., Patterson, A.D., and Benkovic, S.J. (2015). Quantitative analysis of purine nucleotides indicates that purinosomes increase de novo purine biosynthesis. J. Biol. Chem. 290, 6705–6713. 10.1074/jbc.M114.628701.

30. Love, N.R., Chen, Y., Ishibashi, S., Kritsiligkou, P., Lea, R., Koh, Y., Gallop, J.L., Dorey, K., and Amaya, E. (2013). Amputation-induced reactive oxygen species are required for successful Xenopus tadpole tail regeneration. Nat. Cell Biol. 15, 222–228. 10.1038/ncb2659.

31. Love, N.R., Ziegler, M., Chen, Y., and Amaya, E. (2014). Carbohydrate metabolism during vertebrate appendage regeneration: what is its role? How is it regulated?: A postulation that regenerating vertebrate appendages facilitate glycolytic and pentose phosphate pathways to fuel macromolecule biosynthesis. BioEssays News Rev. Mol. Cell. Dev. Biol. 36, 27–33. 10.1002/bies.201300110.

32. Khokha, M.K., Chung, C., Bustamante, E.L., Gaw, L.W.K., Trott, K.A., Yeh, J., Lim, N., Lin, J.C.Y., Taverner, N., Amaya, E., et al. (2002). Techniques and probes for the study of Xenopus tropicalis development. Dev. Dyn. Off. Publ. Am. Assoc. Anat. 225, 499–510. 10.1002/dvdy.10184.

33. Sive, H.L., Grainger, R.M., and Harland, R.M. (2000). Early Development of Xenopus Laevis: A Laboratory Manual (CSHL Press).

34. Nieuwkoop, P.D., and Fader, J. (1994). Normal table of Xenopus laevis (Daudin): a systematical and chronological survey of the development from the fertilized egg till the end of metamorphosis.

35. Schindelin, J., Arganda-Carreras, I., Frise, E., Kaynig, V., Longair, M., Pietzsch, T., Preibisch, S., Rueden, C., Saalfeld, S., Schmid, B., et al. (2012). Fiji: an open-source platform for biological-image analysis. Nat. Methods 9, 676–682. 10.1038/nmeth.2019.

36. Kieserman, E.K., Lee, C., Gray, R.S., Park, T.J., and Wallingford, J.B. (2010). High-magnification in vivo imaging of Xenopus embryos for cell and developmental biology. Cold Spring Harb. Protoc. 2010, pdb.prot5427. 10.1101/pdb.prot5427.

37. Wickham, H. (2009). ggplot2: Elegant Graphics for Data Analysis (Springer) 10.1007/978-0-387-98141-3.

38. Robinson, M.D., McCarthy, D.J., and Smyth, G.K. (2010). edgeR: a Bioconductor package for differential expression analysis of digital gene expression data. Bioinforma. Oxf. Engl. 26, 139–140. 10.1093/bioinformatics/btp616.

39. R Core Team (2024). R: A Language and Environment for Statistical Computing. Version 4.4.1 (R Foundation for Statistical Computing).

