## Supplemental Figures for "Appendage regeneration requires IMPDH2 and creates a sensitized environment for enzyme filament formation"

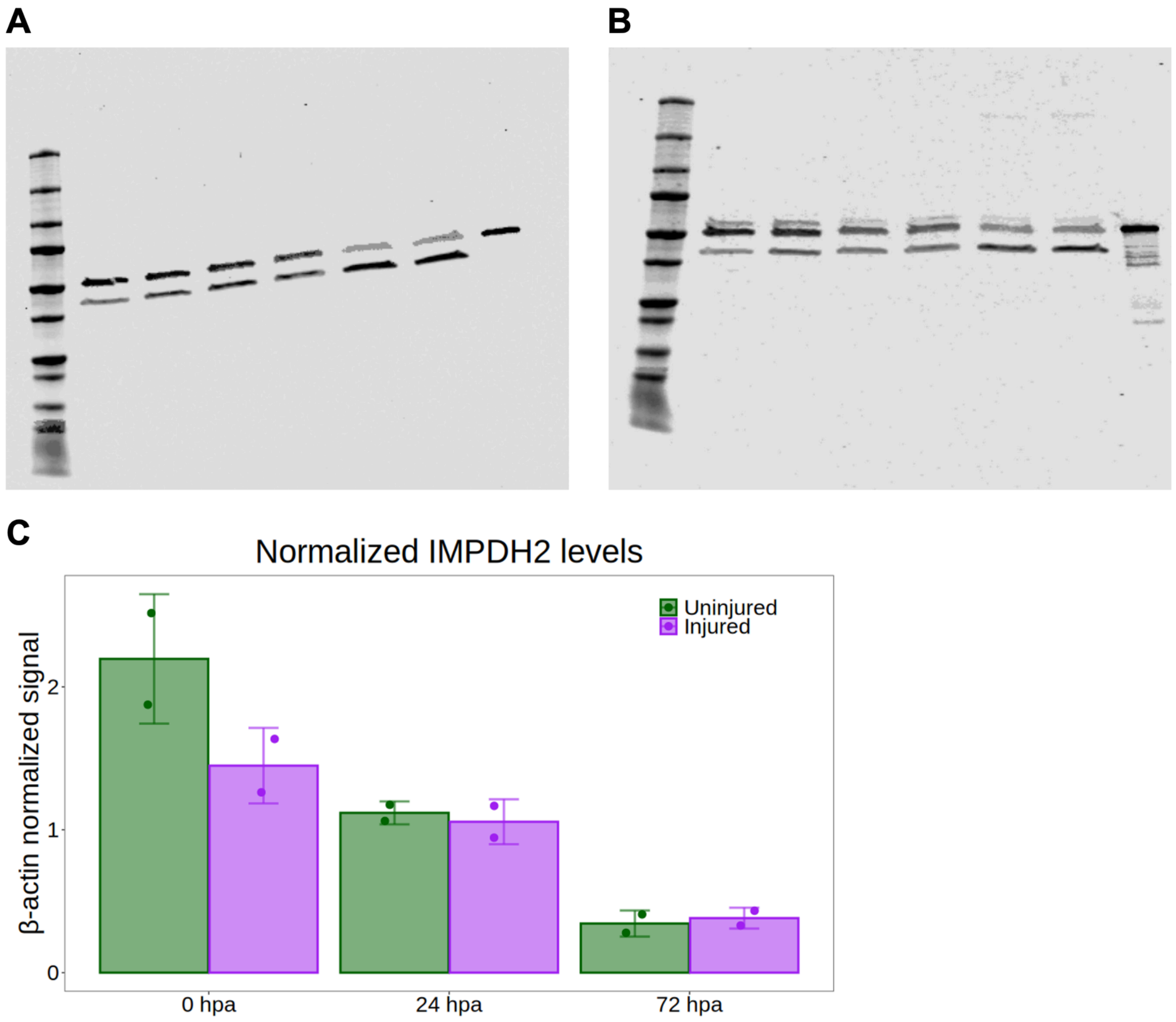


**Supplementary Figure 1. Western blot images and analysis**, related to Figure 2.

(A-B) Uncropped images of western blots. Left to right in each image: ladder, uninjured 0 hpa, injured 0 hpa, uninjured 24 hpa, injured 24 hpa, uninjured 72 hpa, injured 72 hpa, purified hIMPDH2.

(C) Quantification of average IMPDH2 signal intensity at each timepoint normalized to β-actin loading control. Error bars represent +/- S.D.

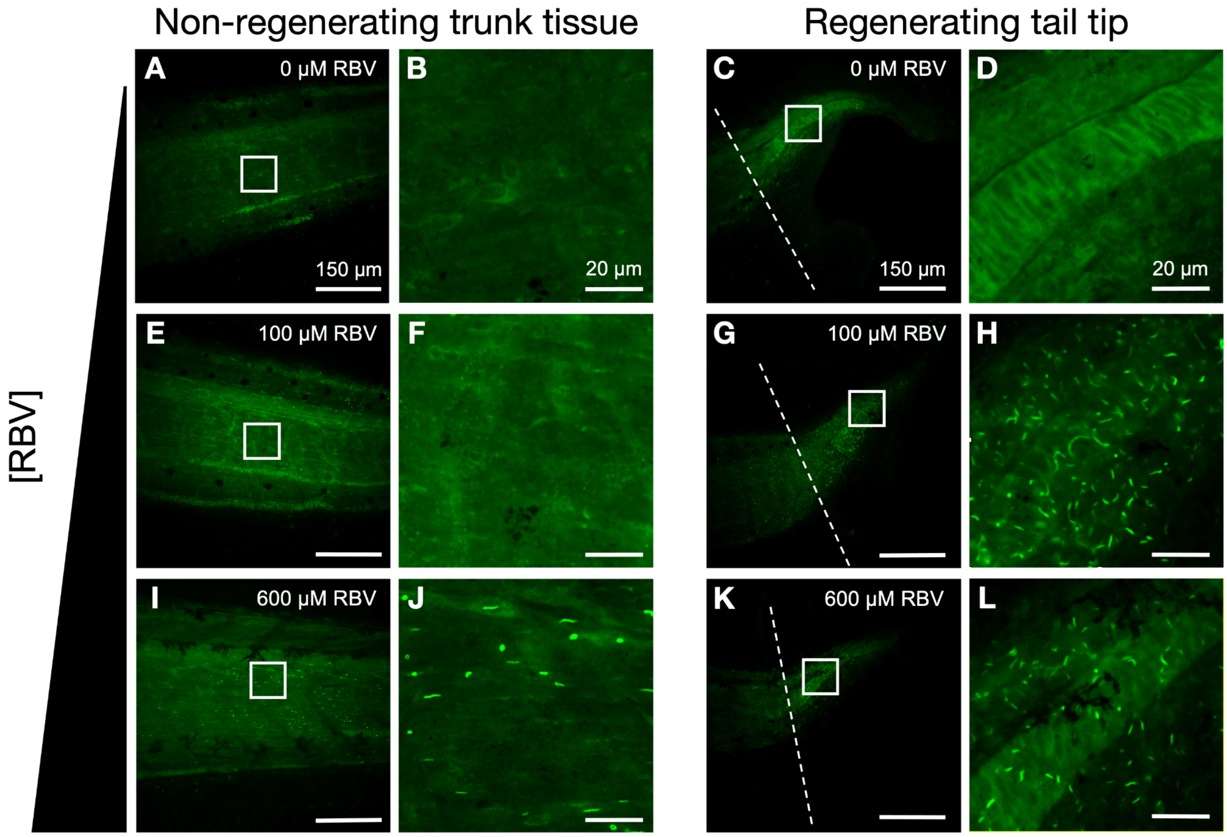


**Supplementary Figure 2. Ribavirin induces superstructures in the sensitized regenerating tail**, related to Figure 4.

(A-D) IHC for IMPDH2 in 72 hpa tadpoles treated with 0.05% DMSO. IMPDH2 filaments rarely assemble under control conditions in the non-regenerating trunk tissue (A and B) or the regenerating tail tip (C and D).

(E-H) IHC for IMPDH2 in 72 hpa tadpoles treated with 100 µM RBV. IMPDH2 filaments do not assemble in the non-regenerating trunk tissue (E and F). Filaments do assemble in the regenerating tail tissue (G and H).

(I-L) IHC for IMPDH2 in 72 hpa tadpoles treated with 600 µM RBV. IMPDH2 forms few filaments in the non-regenerating trunk tissue (I and J). IMPDH2 filaments assemble robustly in the regenerating tail tip (K and L).

Dashed lines in (C), (G), and (K) mark amputation.
